## Supplementary material for "CBP-IDRs regulate acetylation and gene expression": Pairwise sequence alignment

### CBP>p300 (full length | CBP Residues 1-2441)

#### Global Alignment - EMBOSS Needle

```
########################################
# Program: needle
# Rundate: Tue  2 Apr 2024 14:12:33
# Commandline: needle
#    -auto
#    -stdout
#    -asequence emboss_needle-I20240402-141222-0147-12332994-p1m.asequence
#    -bsequence emboss_needle-I20240402-141222-0147-12332994-p1m.bsequence
#    -datafile EBLOSUM62
#    -gapopen 10.0
#    -gapextend 0.5
#    -endopen 10.0
#    -endextend 0.5
#    -aformat3 pair
#    -sprotein1
#    -sprotein2
# Align_format: pair
# Report_file: stdout
########################################

#=======================================
#
# Aligned_sequences: 2
# 1: Hs_crebbp_protein
# 2: EP300_HUMAN
# Matrix: EBLOSUM62
# Gap_penalty: 10.0
# Extend_penalty: 0.5
#
# Length: 2576
# Identity:    1540/2576 (59.8%)
# Similarity:  1795/2576 (69.7%)
# Gaps:         296/2576 (11.5%)
# Score: 7466.5
#
#
#=======================================

Hs_crebbp_pro      1 MAENLLD-GPPNPKRAKLSSPGF--SANDSTDFGSLFDLENDLPDELIPN     47
                     ||||::: |||:.||.|||||..  ||:|.||||||||||:||||||| |
EP300_HUMAN        1 MAENVVEPGPPSAKRPKLSSPALSASASDGTDFGSLFDLEHDLPDELI-N     49

Hs_crebbp_pro     48 GGELGLLNSGN---------LVPDAASKHKQLSELLRGGSGSSINPGI--     86
                     ..||||.|.|:         :|.||||||||||||||.||..::|.|:
EP300_HUMAN       50 STELGLTNGGDINQLQTSLGMVQDAASKHKQLSELLRSGSSPNLNMGVGG     99

Hs_crebbp_pro     87 -GNVSASSPVQQGLGGQAQGQPNSANMASLSAMGKSPLSQGDSSAPSLPK    135
                      |.|.||           |.|.:|..:..:::|.|||::|...::|::
EP300_HUMAN      100 PGQVMAS-----------QAQQSSPGLGLINSMVKSPMTQAGLTSPNM--    136

Hs_crebbp_pro    136 QAASTSGPTPAASQAL----NPQAQKQVGLATSSPATSQTGPGICMNANF    181
                      ...||||....:|:.    :|..|..:|:.|.            |||..
```

```
EP300_HUMAN       137 -GMGTSGPNQGPTQSTGMMNSPVNQPAMGMNTG------------MNAGM     173

Hs_crebbp_pro     182 NQTHPGLLNSNSGHSLINQASQGQAQVMNGSLGAAGRGRGAGMPYPTPAM     231
                      |   ||:|.:.:|..::..      ||||||:| ||||| ..|.||.|.|
EP300_HUMAN       174 N---PGMLAAGNGQGIMPN------QVMNGSIG-AGRGR-QNMQYPNPGM     212

Hs_crebbp_pro     232 QGASSSVLAETLTQVSPQMTGHAGLNTAQAGGMAKMGITGNTSPFGQPFS     281
                       |::.::|.|.|.|.||||.|..||...|.   .|||:..|.:|:|.|::
EP300_HUMAN       213 -GSAGNLLTEPLQQGSPQMGGQTGLRGPQP---LKMGMMNNPNPYGSPYT     258

Hs_crebbp_pro     282 QAGGQPMGATGVNPQLASKQSMVNSLPTFPTDIKNTSVTNVPNMSQMQT-     330
                      |..||.:||:|:..|:.:|..:.|:|..|..|.|......:|||.|...
EP300_HUMAN       259 QNPGQQIGASGLGLQIQTKTVLSNNLSPFAMDKKAVPGGGMPNMGQQPAP     308

Hs_crebbp_pro     331 ---SVGIVP--TQAIATGP-TADPEKRKLIQQQLVLLLHAHKCQRREQAN     374
                         ..|:|.  .|.:.:|. ||||||||||||||||||||||||||||
EP300_HUMAN       309 QVQQPGLVTPVAQGMGSGAHTADPEKRKLIQQQLVLLLHAHKCQRREQAN     358

Hs_crebbp_pro     375 GEVRACSLPHCRTMKNVLNHMTHCQAGKACQVAHCASSRQIISHWKNCTR     424
                      ||||.|:|||||||||||||||||:||:||||||||||||||||||||||
EP300_HUMAN       359 GEVRQCNLPHCRTMKNVLNHMTHCQSGKSCQVAHCASSRQIISHWKNCTR     408

Hs_crebbp_pro     425 HDCPVCLPLKNASDKRNQQTILGSPASGIQNTIGSVGTGQQNATSLSNPN     474
                      |||||||||||.||||||.||.....|:.|. .|:|.|||:|.:|||..:
EP300_HUMAN       409 HDCPVCLPLKNAGDKRNQQPILTGAPVGLGNP-SSLGVGQQSAPNLSTVS     457

Hs_crebbp_pro     475 PIDPSSMQRAYAALGLPY-MNQPQTQLQPQVPGQQPAQP-QTHQQMRTLN     522
                      .|||||::|||||||||| :||..||.|.|...||..|| |:.|.||.::
EP300_HUMAN       458 QIDPSSIERAYAALGLPYQVNQMPTQPQVQAKNQQNQQPGQSPQGMRPMS     507

Hs_crebbp_pro     523 PLGNNPMNIPAG-GITTDQQPPNLISESALPTSLGATNPLMNDGSNSGNI     571
                      .:..:||.:..| |:    |.|:|:|:|.|.:::.:.||:|   |.:.::
EP300_HUMAN       508 NMSASPMGVNGGVGV----QTPSLLSDSMLHSAINSQNPMM---SENASV     550

Hs_crebbp_pro     572 GTLSTIPTAAPPSSTGVRKGWHEHVTQDLRSHLVHKLVQAIFPTPDPAAL     621
                      .:|..:||||.||:||:||.|||.:|||||.||||||||||||||||||
EP300_HUMAN       551 PSLGPMPTAAQPSTTGIRKQWHEDITQDLRNHLVHKLVQAIFPTPDPAAL     600

Hs_crebbp_pro     622 KDRRMENLVAYAKKVEGDMYESANSRDEYYHLLAEKIYKIQKELEEKRRS     671
                      ||||||||||||:||||||||||:|.|||||||||||||||||||||||:
EP300_HUMAN       601 KDRRMENLVAYARKVEGDMYESANNRAEYYHLLAEKIYKIQKELEEKRRT     650

Hs_crebbp_pro     672 RLHKQGILGNQPALPAPGAQP-PVIPQAQPVRPPNGPL--------SLP-     711
                      ||.||.:|.|...:......| |.:.|.||....||||        |:|
EP300_HUMAN       651 RLQKQNMLPNAAGMVPVSMNPGPNMGQPQPGMTSNGPLPDPSMIRGSVPN     700

Hs_crebbp_pro     712 --VNRMQVSQGMNSFNPMSLGNVQLPQAPMGPRAASPMNHSVQMNSMGSV     759
                        :.|:....|:|.|..||:.     |.|:.||...|:.|..|:...|::
EP300_HUMAN       701 QMMPRITPQSGLNQFGQMSMA-----QPPIVPRQTPPLQHHGQLAQPGAL     745

Hs_crebbp_pro     760 -PGMAISPSRMPQPPNMMGAHTNNMMAQAPAQSQFLPQNQFPSSS-GAMS     807
                       |.|...| ||.||.|               |.|||||.||||.. ...:
EP300_HUMAN       746 NPPMGYGP-RMQQPSN---------------QGQFLPQTQFPSQGMNVTN     779

Hs_crebbp_pro     808 VGMGQPPAQTGVSQGQVPGAALP--NPLNMLGPQASQLPCPPVTQSPLH-     854
                      :.:.....|..|||.|:..::.|  :|:...|.|.|.|.:.||.:.|..||
EP300_HUMAN       780 IPLAPSSGQAPVSQAQMSSSSCPVNSPIMPPGSQGSHIHCPQLPQPALHQ     829

Hs_crebbp_pro     855 --PTPPPASTAAGMPSLQHTTPPGMTPPQPAAPTQPSTPVSSSGQTPTPT     902
                        |:|.|:.|..     .|.|||.:...||.|.|.|:          |.||
```

```
EP300_HUMAN      830 NSPSPVPSRTPT-----PHHTPPSIGAQQPPATTIPA---------PVPT    865

Hs_crebbp_pro    903 PGSVPSATQTQS-------TPTVQAAAQAQVTPQPQTP--VQPPSVATPQ    943
                     |.::|...|:|:       |||         .|..|.|  |||...|.|.
EP300_HUMAN      866 PPAMPPGPQSQALHPPPRQTPT---------PPTTQLPQQVQPSLPAAPS    906

Hs_crebbp_pro    944 SS--QQQP--------------TPVHAQPPGTPLSQAAASIDNRVPTPSS    977
                     :.  ||||              .|:....|.|||||.|.||:.:|..|.|
EP300_HUMAN      907 ADQPQQQPRSQQSTAASVPTPTAPLLPPQPATPLSQPAVSIEGQVSNPPS    956

Hs_crebbp_pro    978 VASAETNSQQPGPDVPVLEMKTETQAEDTEPDPGESKGEPRSEMMEEDLQ   1027
                     .:|.|.|||......|..|:|.|.:.|..:|:|.:::.|..||...||.
EP300_HUMAN      957 TSSTEVNSQAIAEKQPSQEVKMEAKMEVDQPEPADTQPEDISESKVEDC-   1005

Hs_crebbp_pro   1028 GASQVKEETDIAEQKSEPMEVDEKKPEVKVEVKEEEESSSNGTASQSTSP   1077
                                   |.|..|.:|:..|:|.|:||||:..|......|.:|
EP300_HUMAN     1006 --------------KMESTETEERSTELKTEIKEEEDQPSTSATQSSPAP   1041

Hs_crebbp_pro   1078 SQPRKKIFKPEELRQALMPTLEALYRQDPESLPFRQPVDPQLLGIPDYFD   1127
                     .|.:||||||||||||||||||||||||||||||||||||||||||||||
EP300_HUMAN     1042 GQSKKKIFKPEELRQALMPTLEALYRQDPESLPFRQPVDPQLLGIPDYFD   1091

Hs_crebbp_pro   1128 IVKNPMDLSTIKRKLDTGQYQEPWQYVDDVWLMFNNAWLYNRKTSRVYKF   1177
                     |||:|||||||||||||||||||||||||:||||||||||||||||||||:
EP300_HUMAN     1092 IVKSPMDLSTIKRKLDTGQYQEPWQYVDDIWLMFNNAWLYNRKTSRVYKY   1141

Hs_crebbp_pro   1178 CSKLAEVFEQEIDPVMQSLGYCCGRKYEFSPQTLCCYGKQLCTIPRDAAY   1227
                     ||||:|||||||||||||||||||||.|||||||||||||||||||||.|
EP300_HUMAN     1142 CSKLSEVFEQEIDPVMQSLGYCCGRKLEFSPQTLCCYGKQLCTIPRDATY   1191

Hs_crebbp_pro   1228 YSYQNRYHFCEKCFTEIQGENVTLGDDPSQPQTTISKDQFEKKKNDTLDP   1277
                     ||||||||||||||.|||||:|:||||||||||||:|:||.|:|||||||
EP300_HUMAN     1192 YSYQNRYHFCEKCFNEIQGESVSLGDDPSQPQTTINKEQFSKRKNDTLDP   1241

Hs_crebbp_pro   1278 EPFVDCKECGRKMHQICVLHYDIIWPSGFVCDNCLKKTGRPRKENKFSAK   1327
                     |.||:|.|||||||||||||::||||:|||||.||||:.|.|||||||||
EP300_HUMAN     1242 ELFVECTECGRKMHQICVLHHEIIWPAGFVCDGCLKKSARTRKENKFSAK   1291

Hs_crebbp_pro   1328 RLQTTRLGNHLEDRVNKFLRRQNHPEAGEVFVRVVASSDKTVEVKPGMKS   1377
                     ||.:||||..||:|||.|||||||||:|||.||||.:||||||||||||:
EP300_HUMAN     1292 RLPSTRLGTFLENRVNDFLRRQNHPESGEVTVRVVHASDKTVEVKPGMKA   1341

Hs_crebbp_pro   1378 RFVDSGEMSESFPYRTKALFAFEEIDGVDVCFFGMHVQEYGSDCPPPNTR   1427
                     ||||||||:|||||||||||||||||||||:||||||||||||||||||.|
EP300_HUMAN     1342 RFVDSGEMAESFPYRTKALFAFEEIDGVDLCFFGMHVQEYGSDCPPPNQR   1391

Hs_crebbp_pro   1428 RVYISYLDSIHFFRPRCLRTAVYHEILIGYLEYVKKLGYVTGHIWACPPS   1477
                     |||||||||:|||||:||||||||||||||||||||||||.||||||||||
EP300_HUMAN     1392 RVYISYLDSVHFFRPKCLRTAVYHEILIGYLEYVKKLGYTTGHIWACPPS   1441

Hs_crebbp_pro   1478 EGDDYIFHCHPPDQKIPKPKRLQEWYKKMLDKAFAERIIHDYKDIFKQAT   1527
                     ||||||||||||||||||||||||||||||||||.:|||:||||||||||||
EP300_HUMAN     1442 EGDDYIFHCHPPDQKIPKPKRLQEWYKKMLDKAVSERIVHDYKDIFKQAT   1491

Hs_crebbp_pro   1528 EDRLTSAKELPYFEGDFWPNVLEESIKELEQEEEERKKEESTAASETTEG   1577
                     ||||||||||||||||||||||||||||||||||||||:||:| ::|:|:.
EP300_HUMAN     1492 EDRLTSAKELPYFEGDFWPNVLEESIKELEQEEEERKREENT-SNESTDV   1540

Hs_crebbp_pro   1578 SQGDSKNAKKKNNKKTNKNKSSISRANKKKPSMPNVSNDLSQKLYATMEK   1627
                     ::|||||||||||||:|||||:||.|||||.||||||||||||||||||
```

```
EP300_HUMAN      1541 TKGDSKNAKKKNNKKTSKNKSSLSRGNKKKPGMPNVSNDLSQKLYATMEK   1590

Hs_crebbp_pro    1628 HKEVFFVIHLHAGPVINTLPPIVDPDPLLSCDLMDGRDAFLTLARDKHWE   1677
                      ||||||||.|.|||..|:||||||||||:.|||||||||||||||||||.|
EP300_HUMAN      1591 HKEVFFVIRLIAGPAANSLPPIVDPDPLIPCDLMDGRDAFLTLARDKHLE   1640

Hs_crebbp_pro    1678 FSSLRRSKWSTLCMLVELHTQGQDRFVYTCNECKHHVETRWHCTVCEDYD   1727
                      ||||||::|||:||||||||.|||||||||||||||||||||||||||||
EP300_HUMAN      1641 FSSLRRAQWSTMCMLVELHTQSQDRFVYTCNECKHHVETRWHCTVCEDYD   1690

Hs_crebbp_pro    1728 LCINCYNTKSHAHKMVKWGLGLDDEGSSQGEPQSKSPQESRRLSIQRCIQ   1777
                      |||.|||||:|.|||.|.|||||||.::|....::||.:|||||||||||
EP300_HUMAN      1691 LCITCYNTKNHDHKMEKLGLGLDDESNNQQAAATQSPGDSRRLSIQRCIQ   1740

Hs_crebbp_pro    1778 SLVHACQCRNANCSLPSCQKMKRVVQHTKGCKRKTNGGCPVCKQLIALCC   1827
                      ||||||||||||||||||||||||||||||||||||||||:|||||||||
EP300_HUMAN      1741 SLVHACQCRNANCSLPSCQKMKRVVQHTKGCKRKTNGGCPICKQLIALCC   1790

Hs_crebbp_pro    1828 YHAKHCQENKCPVPFCLNIKHKLRQQQIQHRLQQAQLMRRRMATMNTRNV   1877
                      |||||||||||||||||||.||||||:||||||||::|||||:|....|
EP300_HUMAN      1791 YHAKHCQENKCPVPFCLNIKQKLRQQQLQHRLQQAQMLRRRMASMQRTGV   1840

Hs_crebbp_pro    1878 --PQQSLPSPTSAPPGTPT-QQPSTPQTPQPPAQPQPSPVSMSPAGFPSV   1924
                        .||.|||||.|.|.||| |||:|||||||.:||||:|.:..|    |.:
EP300_HUMAN      1841 VGQQQGLPSPTPATPTTPTGQQPTTPQTPQPTSQPQPTPPNSMP---PYL   1887

Hs_crebbp_pro    1925 ARTQPPTTVSTGKPTSQV--PAPPPPAQ-----PPPAAVEAARQIEREAQ   1967
                      .|||....||.||...||  |.||..||     ||||||.|.||:|.|:
EP300_HUMAN      1888 PRTQAAGPVSQGKAAGQVTPPTPPQTAQPPLPGPPPAAVEMAMQIQRAAE   1937

Hs_crebbp_pro    1968 QQQHLYRVN-----INNSMPPGRTGMGTPGSQMAPVSLNVPRPNQ-VSGP   2011
                      .|:.:..|.     |.:.|||     .||   |||:.:|.|...: .||.
EP300_HUMAN      1938 TQRQMAHVQIFQRPIQHQMPP-----MTP---MAPMGMNPPPMTRGPSGH   1979

Hs_crebbp_pro    2012 VMPSM-PPGQ-----WQQAPLPQ-QQPMPGLPRPVISMQAQ-------AA   2047
                      :.|.| |.|.     |.|..||| ||...|:|||.:...||       |.
EP300_HUMAN      1980 LEPGMGPTGMQQQPPWSQGGLPQPQQLQSGMPRPAMMSVAQHGQPLNMAP   2029

Hs_crebbp_pro    2048 VAGPRMPSVQP--PRSISPSALQDLLRTLKSPSSPQQQQQVLNILKSNPQ   2095
                      ..|.....:.|  |.::|..|||:|||||:|||||.||||||:||.:|||
EP300_HUMAN      2030 QPGLGQVGISPLKPGTVSQQALQNLLRTLRSPSSPLQQQQVLSILHANPQ   2079

Hs_crebbp_pro    2096 LMAAFIKQRTAKYVANQPGMQP---QPGL-QSQPGMQP-----QPGMHQQ   2136
                      |:|||||||.|||..:.|  ||   |||: |.|||:||     |.|:|..
EP300_HUMAN      2080 LLAAFIKQRAAKYANSNP--QPIPGQPGMPQGQPGLQPPTMPGQQGVHSN   2127

Hs_crebbp_pro    2137 PSLQNLNAMQAGVPRPGVP---PQQQ---AMGGLNPQGQALNIMNPGHNP   2180
                      |::||:|.|||||.|.|:|   ||||   .|||::||.|.:| ||  ||
EP300_HUMAN      2128 PAMQNMNPMQAGVQRAGLPQQQPQQQLQPPMGGMSPQAQQMN-MN--HN-   2173

Hs_crebbp_pro    2181 NMASMNPQYREMLRRQLLQQQQQQQQQQQQQQQQQGSAGMAGGMAGHGQ   2230
                         :|..|:|::|||          ||..||||||....|:..|||.|.|
EP300_HUMAN      2174 ---TMPSQFRDILRR----------QQMMQQQQQQGAGPGIGPGMANHNQ   2210

Hs_crebbp_pro    2231 FQQPQGPGGYPPAMQQQQRMQQHLPLQGSSMGQMAAQMGQLGQMGQ--PG   2278
                      ||||||. ||||  |||||||.|       |.||  |.|.:||:||  ..
EP300_HUMAN      2211 FQQPQGV-GYPP--QQQQRMQHH-------MQQM--QQGNMGQIGQLPQA   2248

Hs_crebbp_pro    2279 LGADSTPNIQQALQQRILQQQQMKQQIGSPGQPNPMSPQQHMLSGQPQAS   2328
                      |||::..:: ||.|||:||     ||:|||.||||||||||||..|.|:.
```

```
EP300_HUMAN     2249 LGAEAGASL-QAYQQRLLQ-----QQMGSPVQPNPMSPQQHMLPNQAQSP   2292

Hs_crebbp_pro   2329 HLPGQQIATSLSNQVRSPAPVQSPRPQSQPPHSSPSPRIQPQPSPHHVSP   2378
                     ||.||||..|||||||||.||.||||||||||||||||:|||||||||||
EP300_HUMAN     2293 HLQGQQIPNSLSNQVRSPQPVPSPRPQSQPPHSSPSPRMQPQPSPHHVSP   2342

Hs_crebbp_pro   2379 QTGSPHPGLAVTMASSIDQGHLGNPEQSAMLPQL-NTPS----RSALSSE   2423
                     ||.||||||....|:.::|||..:|:|::||.|| :.|.    ..|.:::
EP300_HUMAN     2343 QTSSPHPGLVAAQANPMEQGHFASPDQNSMLSQLASNPGMANLHGASATD   2392

Hs_crebbp_pro   2424 LSLVGDTTG-------DTLEKFVEGL   2442
                     |.|..|.:.       .||:..
EP300_HUMAN     2393 LGLSTDNSDLNSNLSQSTLDIH----   2414


#---------------------------------------
#---------------------------------------
```

### CBP>p300 (full length | CBP Residues 1-2441)

#### Local Alignment - EMBOSS Matcher

```
########################################
# Program: matcher
# Rundate: Tue  2 Apr 2024 14:02:58
# Commandline: matcher
#    -auto
#    -stdout
#    -asequence emboss_matcher-I20240402-140245-0831-23478561-p1m.asequence
#    -bsequence emboss_matcher-I20240402-140245-0831-23478561-p1m.bsequence
#    -datafile EBLOSUM62
#    -gapopen 14
#    -gapextend 4
#    -alternatives 1
#    -aformat3 pair
#    -sprotein1
#    -sprotein2
# Align_format: pair
# Report_file: stdout
########################################

#=======================================
#
# Aligned_sequences: 2
# 1: Hs_crebbp_protein
# 2: EP300_HUMAN
# Matrix: EBLOSUM62
# Gap_penalty: 14
# Extend_penalty: 4
#
# Length: 2485
# Identity:    1512/2485 (60.8%)
# Similarity:  1758/2485 (70.7%)
# Gaps:         182/2485 ( 7.3%)
# Score: 6735
#
#
#=======================================

Hs_crebbp_pro      1 MAENLLD-GPPNPKRAKLSSPGFSAN--DSTDFGSLFDLENDLPDELIPN     47
                     ||||::: |||:.||.|||||..||:  |.||||||||||:||||||| |
EP300_HUMAN        1 MAENVVEPGPPSAKRPKLSSPALSASASDGTDFGSLFDLEHDLPDELI-N     49

Hs_crebbp_pro     48 GGELGLLNSGNL---------VPDAASKHKQLSELLRGGSGSSINPGIGN     88
                     ..||||.|.|::         |.||||||||||||||.||..::|.|:|.
EP300_HUMAN       50 STELGLTNGGDINQLQTSLGMVQDAASKHKQLSELLRSGSSPNLNMGVGG     99

Hs_crebbp_pro     89 VSASSPVQQGLGGQAQGQPNSANMASLSAMGKSPLSQGDSSAPSLPKQAA    138
                             .|....:|.|.:|..:..:::|.|||::|...::|::   ..
EP300_HUMAN      100 --------PGQVMASQAQQSSPGLGLINSMVKSPMTQAGLTSPNM---GM    138

Hs_crebbp_pro    139 STSGPTPAASQALNPQAQKQVGLATSSPATSQTGPGICMNANFNQTHPGL    188
                     .||||....:|:        .|:..|.......|....|||..|   ||:
EP300_HUMAN      139 GTSGPNQGPTQS--------TGMMNSPVNQPAMGMNTGMNAGMN---PGM    177

Hs_crebbp_pro    189 LNSNSGHSLINQASQGQAQVMNGSLGAAGRGRGAGMPYPTPAMQGASSSV    238
```

```
                          |.:.:|..::..      ||||||:|| |||| ..|.||.|.| |::.::
EP300_HUMAN       178 LAAGNGQGIMPN------QVMNGSIGA-GRGR-QNMQYPNPGM-GSAGNL     218

Hs_crebbp_pro     239 LAETLTQVSPQMTGHAGLNTAQAGGMAKMGITGNTSPFGQPFSQAGGQPM    288
                      |.|.|.|.||||.|..||...|.    .|||:..|.:|:|.|::|..||.:
EP300_HUMAN       219 LTEPLQQGSPQMGGQTGLRGPQP---LKMGMMNNPNPYGSPYTQNPGQQI    265

Hs_crebbp_pro     289 GATGVNPQLASKQSMVNSLPTFPTDIKNTSVTNVPNMSQMQT----SVGI    334
                      ||:|:..|:.:|..:.|:|..|..|.|......:|||.|...    ..|:
EP300_HUMAN       266 GASGLGLQIQTKTVLSNNLSPFAMDKKAVPGGGMPNMGQQPAPQVQQPGL    315

Hs_crebbp_pro     335 VPTQAIATGP---TADPEKRKLIQQQLVLLLHAHKCQRREQANGEVRACS    381
                      |...|...|.   |||||||||||||||||||||||||||||||||.|:
EP300_HUMAN       316 VTPVAQGMGSGAHTADPEKRKLIQQQLVLLLHAHKCQRREQANGEVRQCN    365

Hs_crebbp_pro     382 LPHCRTMKNVLNHMTHCQAGKACQVAHCASSRQIISHWKNCTRHDCPVCL    431
                      |||||||||||||||||:||:|||||||||||||||||||||||||||||
EP300_HUMAN       366 LPHCRTMKNVLNHMTHCQSGKSCQVAHCASSRQIISHWKNCTRHDCPVCL    415

Hs_crebbp_pro     432 PLKNASDKRNQQTILGSPASGIQNTIGSVGTGQQNATSLSNPNPIDPSSM    481
                      |||||.||||||.||.....|:.|. .|:|.|||:|.:||..:.|||||:
EP300_HUMAN       416 PLKNAGDKRNQQPILTGAPVGLGNP-SSLGVGQQSAPNLSTVSQIDPSSI    464

Hs_crebbp_pro     482 QRAYAALGLPY-MNQPQTQLQPQVPGQQPAQP-QTHQQMRTLNPLGNNPM    529
                      :|||||||||| :||..||.|.|...||..|| |:.|.||.::.:..:||
EP300_HUMAN       465 ERAYAALGLPYQVNQMPTQPQVQAKNQQNQQPGQSPQGMRPMSNMSASPM    514

Hs_crebbp_pro     530 NIPAGGITTDQQPPNLISESALPTSLGATNPLMNDGSNSGNIGTLSTIPT    579
                      .:. ||:..   |.|:|:|:|.|.:::.:.||:|   |.:.::.:|..:||
EP300_HUMAN       515 GVN-GGVGV--QTPSLLSDSMLHSAINSQNPMM---SENASVPSLGPMPT    558

Hs_crebbp_pro     580 AAPPSSTGVRKGWHEHVTQDLRSHLVHKLVQAIFPTPDPAALKDRRMENL    629
                      ||.||:||:||.|||.:|||||:|||||||||||||||||||||||||||
EP300_HUMAN       559 AAQPSTTGIRKQWHEDITQDLRNHLVHKLVQAIFPTPDPAALKDRRMENL    608

Hs_crebbp_pro     630 VAYAKKVEGDMYESANSRDEYYHLLAEKIYKIQKELEEKRRSRLHKQGIL    679
                      ||||:||||||||||||:|.||||||||||||||||||||||:||.||.:|
EP300_HUMAN       609 VAYARKVEGDMYESANNRAEYYHLLAEKIYKIQKELEEKRRTRLQKQNML    658

Hs_crebbp_pro     680 GNQPALPAPGAQP-PVIPQAQPVRPPNGPLSLP--VNRMQVSQGMNSFNP    726
                      .|...:......| |.:.|.||....||||..|  :.....:|.|.....|
EP300_HUMAN       659 PNAAGMVPVSMNPGPNMGQPQPGMTSNGPLPDPSMIRGSVPNQMMPRITP    708

Hs_crebbp_pro     727 MS----LGNVQLPQAPMGPRAASPMNHSVQMNSMGSV-PGMAISPSRMPQ    771
                      .|    .|.:.:.|.|:.||...|:.|..|:...|:: |.|...| ||.|
EP300_HUMAN       709 QSGLNQFGQMSMAQPPIVPRQTPPLQHHGQLAQPGALNPPMGYGP-RMQQ    757

Hs_crebbp_pro     772 PPNMMGAHTNNMMAQAPAQSQFLPQNQFPSSS-GAMSVGMGQPPAQTGVS    820
                      |.|               |.|||||.||||.. ...::.:.....|..||
EP300_HUMAN       758 PSN---------------QGQFLPQTQFPSQGMNVTNIPLAPSSGQAPVS    792

Hs_crebbp_pro     821 QGQVPGAALP--NPLNMLGPQASQLPCPPVTQSPLH---PTPPPASTAAG    865
                      |.|:..::.|  :|:...|.|.|.:.||.:.:|..||   |:|.|:.|...
EP300_HUMAN       793 QAQMSSSSCPVNSPIMPPGSQGSHIHCPQLPQPALHQNSPSPVPSRTPTP    842

Hs_crebbp_pro     866 M---PSLQHTTPPGMTPPQPAAPTQPSTPVSSSGQTPTPTPGSVPSATQT    912
                      .   ||:....||..|.|.|. ||.|:.|.....|...|.|...|:...|
EP300_HUMAN       843 HHTPPSIGAQQPPATTIPAPV-PTPPAMPPGPQSQALHPPPRQTPTPPTT    891

Hs_crebbp_pro     913 QSTPTVQAAAQAQVTPQPQTPVQPPSVATPQSSQQQPT-PVHAQPPGTPL    961
```

```
                      |....||.:..|..:.. |...||.|..:..:|...|| |:....|.|||
EP300_HUMAN       892 QLPQQVQPSLPAAPSAD-QPQQQPRSQQSTAASVPTPTAPLLPPQPATPL     940

Hs_crebbp_pro     962 SQAAASIDNRVPTPSSVASAETNSQQPGPDVPVLEMKTETQAEDTEPDPG    1011
                      ||.|.||:.:|..|.|.:|.|.|||......|..|:|.|.:.|..:|:|.
EP300_HUMAN       941 SQPAVSIEGQVSNPPSTSSTEVNSQAIAEKQPSQEVKMEAKMEVDQPEPA     990

Hs_crebbp_pro    1012 ESKGEPRSEMMEEDLQGASQVKEETDIAEQKSEPMEVDEKKPEVKVEVKE    1061
                      :::.|..||...||.               |.|..|.:|:..|:|.|:||
EP300_HUMAN       991 DTQPEDISESKVEDC---------------KMESTETEERSTELKTEIKE    1025

Hs_crebbp_pro    1062 EEESSSNGTASQSTSPSQPRKKIFKPEELRQALMPTLEALYRQDPESLPF    1111
                      ||:..|......|.:|.|.:||||||||||||||||||||||||||||||
EP300_HUMAN      1026 EEDQPSTSATQSSPAPGQSKKKIFKPEELRQALMPTLEALYRQDPESLPF    1075

Hs_crebbp_pro    1112 RQPVDPQLLGIPDYFDIVKNPMDLSTIKRKLDTGQYQEPWQYVDDVWLMF    1161
                      |||||||||||||||||||:|||||||||||||||||||||||||:||||
EP300_HUMAN      1076 RQPVDPQLLGIPDYFDIVKSPMDLSTIKRKLDTGQYQEPWQYVDDIWLMF    1125

Hs_crebbp_pro    1162 NNAWLYNRKTSRVYKFCSKLAEVFEQEIDPVMQSLGYCCGRKYEFSPQTL    1211
                      |||||||||||||||:||||:|||||||||||||||||||||.|||||||
EP300_HUMAN      1126 NNAWLYNRKTSRVYKYCSKLSEVFEQEIDPVMQSLGYCCGRKLEFSPQTL    1175

Hs_crebbp_pro    1212 CCYGKQLCTIPRDAAYYSYQNRYHFCEKCFTEIQGENVTLGDDPSQPQTT    1261
                      ||||||||||||||.||||||||||||||||.|||||:|:||||||||||
EP300_HUMAN      1176 CCYGKQLCTIPRDATYYSYQNRYHFCEKCFNEIQGESVSLGDDPSQPQTT    1225

Hs_crebbp_pro    1262 ISKDQFEKKKNDTLDPEPFVDCKECGRKMHQICVLHYDIIWPSGFVCDNC    1311
                      |:|:||.|:||||||||.||:|.|||||||||||||::||||:|||||.|
EP300_HUMAN      1226 INKEQFSKRKNDTLDPELFVECTECGRKMHQICVLHHEIIWPAGFVCDGC    1275

Hs_crebbp_pro    1312 LKKTGRPRKENKFSAKRLQTTRLGNHLEDRVNKFLRRQNHPEAGEVFVRV    1361
                      |||:.|.|||||||||||.:||||..||:|||.|||||||||:|||.|||
EP300_HUMAN      1276 LKKSARTRKENKFSAKRLPSTRLGTFLENRVNDFLRRQNHPESGEVTVRV    1325

Hs_crebbp_pro    1362 VASSDKTVEVKPGMKSRFVDSGEMSESFPYRTKALFAFEEIDGVDVCFFG    1411
                      |.:||||||||||||:||||||||:||||||||||||||||||||:||||
EP300_HUMAN      1326 VHASDKTVEVKPGMKARFVDSGEMAESFPYRTKALFAFEEIDGVDLCFFG    1375

Hs_crebbp_pro    1412 MHVQEYGSDCPPPNTRRVYISYLDSIHFFRPRCLRTAVYHEILIGYLEYV    1461
                      ||||||||||||||.||||||||||:|||||:||||||||||||||||||
EP300_HUMAN      1376 MHVQEYGSDCPPPNQRRVYISYLDSVHFFRPKCLRTAVYHEILIGYLEYV    1425

Hs_crebbp_pro    1462 KKLGYVTGHIWACPPSEGDDYIFHCHPPDQKIPKPKRLQEWYKKMLDKAF    1511
                      |||||.|||||||||||||||||||||||||||||||||||||||||||.
EP300_HUMAN      1426 KKLGYTTGHIWACPPSEGDDYIFHCHPPDQKIPKPKRLQEWYKKMLDKAV    1475

Hs_crebbp_pro    1512 AERIIHDYKDIFKQATEDRLTSAKELPYFEGDFWPNVLEESIKELEQEEE    1561
                      :|||:|||||||||||||||||||||||||||||||||||||||||||||
EP300_HUMAN      1476 SERIVHDYKDIFKQATEDRLTSAKELPYFEGDFWPNVLEESIKELEQEEE    1525

Hs_crebbp_pro    1562 ERKKEESTAASETTEGSQGDSKNAKKKNNKKTNKNKSSISRANKKKPSMP    1611
                      |||:||:|: :|:|:.::||||||||||||||:|||||:||.|||||.||
EP300_HUMAN      1526 ERKREENTS-NESTDVTKGDSKNAKKKNNKKTSKNKSSLSRGNKKKPGMP    1574

Hs_crebbp_pro    1612 NVSNDLSQKLYATMEKHKEVFFVIHLHAGPVINTLPPIVDPDPLLSCDLM    1661
                      ||||||||||||||||||||||||.|.|||..|:||||||||||:.||||
EP300_HUMAN      1575 NVSNDLSQKLYATMEKHKEVFFVIRLIAGPAANSLPPIVDPDPLIPCDLM    1624

Hs_crebbp_pro    1662 DGRDAFLTLARDKHWEFSSLRRSKWSTLCMLVELHTQGQDRFVYTCNECK    1711
```

```
                       ||||||||||||||.|||||||::|||:|||||||||.||||||||||||
EP300_HUMAN       1625 DGRDAFLTLARDKHLEFSSLRRAQWSTMCMLVELHTQSQDRFVYTCNECK   1674

Hs_crebbp_pro     1712 HHVETRWHCTVCEDYDLCINCYNTKSHAHKMVKWGLGLDDEGSSQGEPQS   1761
                       |||||||||||||||||||.|||||:|.|||.|.|||||||.::|....:
EP300_HUMAN       1675 HHVETRWHCTVCEDYDLCITCYNTKNHDHKMEKLGLGLDDESNNQQAAAT   1724

Hs_crebbp_pro     1762 KSPQESRRLSIQRCIQSLVHACQCRNANCSLPSCQKMKRVVQHTKGCKRK   1811
                       :||.:|||||||||||||||||||||||||||||||||||||||||||||
EP300_HUMAN       1725 QSPGDSRRLSIQRCIQSLVHACQCRNANCSLPSCQKMKRVVQHTKGCKRK   1774

Hs_crebbp_pro     1812 TNGGCPVCKQLIALCCYHAKHCQENKCPVPFCLNIKHKLRQQQIQHRLQQ   1861
                       ||||||:|||||||||||||||||||||||||||||.||||||:||||||
EP300_HUMAN       1775 TNGGCPICKQLIALCCYHAKHCQENKCPVPFCLNIKQKLRQQQLQHRLQQ   1824

Hs_crebbp_pro     1862 AQLMRRRMATMNTRNV--PQQSLPSPTSAPPGTPT-QQPSTPQTPQPPAQ   1908
                       ||::|||||:|....|  .||.|||||.|.|.||| |||:|||||||.:|
EP300_HUMAN       1825 AQMLRRRMASMQRTGVVGQQQGLPSPTPATPTTPTGQQPTTPQTPQPTSQ   1874

Hs_crebbp_pro     1909 PQPSPVSMSPAGFPSVARTQPPTTVSTGKPTSQV--PAPPPPAQPP----   1952
                       |||:|.:..|   |.:.|||....||.||...||  |.||..||||
EP300_HUMAN       1875 PQPTPPNSMP---PYLPRTQAAGPVSQGKAAGQVTPPTPPQTAQPPLPGP   1921

Hs_crebbp_pro     1953 -PAAVEAARQIEREAQQQQHLYRVNINNSMPPGRTGMGTPGSQMAPVSLN   2001
                        |||||.|.||:|.|:.|:.:..|.|  ...|.:..| .|.:.|||:.:|
EP300_HUMAN       1922 PPAAVEMAMQIQRAAETQRQMAHVQI--FQRPIQHQM-PPMTPMAPMGMN   1968

Hs_crebbp_pro     2002 VPRPNQ-VSGPVMPSMPPGQWQQAP------LPQ-QQPMPGLPRPVISMQ   2043
                       .|...: .||.:.|.|.|...||.|      ||| ||...|:|||.:...
EP300_HUMAN       1969 PPPMTRGPSGHLEPGMGPTGMQQQPPWSQGGLPQPQQLQSGMPRPAMMSV   2018

Hs_crebbp_pro     2044 AQ-------AAVAGPRMPSVQP--PRSISPSALQDLLRTLKSPSSPQQQQ   2084
                       ||       |...|.....:.|  |.::|..|||:|||||:|||||.|||
EP300_HUMAN       2019 AQHGQPLNMAPQPGLGQVGISPLKPGTVSQQALQNLLRTLRSPSSPLQQQ   2068

Hs_crebbp_pro     2085 QVLNILKSNPQLMAAFIKQRTAKYVANQPGMQP-QPGL-QSQPGMQP---   2129
                       |||:||.:||||:|||||||.|||..:.|...| |||: |.|||:||
EP300_HUMAN       2069 QVLSILHANPQLLAAFIKQRAAKYANSNPQPIPGQPGMPQGQPGLQPPTM   2118

Hs_crebbp_pro     2130 --QPGMHQQPSLQNLNAMQAGVPRPGVPPQQQAMGGLNPQGQALNIMNPG   2177
                         |.|:|..|::||:|.|||||.|.|:| |||....|.|....::.....
EP300_HUMAN       2119 PGQQGVHSNPAMQNMNPMQAGVQRAGLP-QQQPQQQLQPPMGGMSPQAQQ   2167

Hs_crebbp_pro     2178 HNPNMASMNPQYREMLRRQLLQQQQQQQQQQQQQQQQQQGSAGMAGGMAG   2227
                       .|.|..:|..|:|::||||          |..||||||....|:..|||.
EP300_HUMAN       2168 MNMNHNTMPSQFRDILRRQ----------QMMQQQQQQGAGPGIGPGMAN   2207

Hs_crebbp_pro     2228 HGQFQQPQGPGGYPPAMQQQQRMQQHLPLQGSSMGQMAAQMGQLGQMGQP   2277
                       |.|||||||.| |||  |||||||.|:    ..|.|  ..|||:||:.|
EP300_HUMAN       2208 HNQFQQPQGVG-YPP--QQQQRMQHHM----QQMQQ--GNMGQIGQLPQ-   2247

Hs_crebbp_pro     2278 GLGADSTPNIQQALQQRILQQQQMKQQIGSPGQPNPMSPQQHMLSGQPQA   2327
                       .|||::..::| |.|||:||||     :|||.||||||||||||..|.|:
EP300_HUMAN       2248 ALGAEAGASLQ-AYQQRLLQQQ-----MGSPVQPNPMSPQQHMLPNQAQS   2291

Hs_crebbp_pro     2328 SHLPGQQIATSLSNQVRSPAPVQSPRPQSQPPHSSPSPRIQPQPSPHHVS   2377
                       .||.||||..|||||||||.||.||||||||||||||||:||||||||||
EP300_HUMAN       2292 PHLQGQQIPNSLSNQVRSPQPVPSPRPQSQPPHSSPSPRMQPQPSPHHVS   2341

Hs_crebbp_pro     2378 PQTGSPHPGLAVTMASSIDQGHLGNPEQSAMLPQL   2412
```

```
                     |||.||||||....|:.::|||..:|:|::||.||
EP300_HUMAN     2342 PQTSSPHPGLVAAQANPMEQGHFASPDQNSMLSQL   2376


#---------------------------------------
#---------------------------------------
```

### CBP>p300 (HAT domain | CBP Residues 1196-1718)

```
########################################
# Program: matcher
# Rundate: Tue  2 Apr 2024 18:44:14
# Commandline: matcher
#    -auto
#    -stdout
#    -asequence emboss_matcher-I20240402-184400-0923-51175319-p1m.asequence
#    -bsequence emboss_matcher-I20240402-184400-0923-51175319-p1m.bsequence
#    -datafile EBLOSUM62
#    -gapopen 14
#    -gapextend 4
#    -alternatives 1
#    -aformat3 pair
#    -sprotein1
#    -sprotein2
# Align_format: pair
# Report_file: stdout
########################################

#=======================================
#
# Aligned_sequences: 2
# 1: CBP
# 2: p300
# Matrix: EBLOSUM62
# Gap_penalty: 14
# Extend_penalty: 4
#
# Length: 503
# Identity:     439/503 (87.3%)
# Similarity:   475/503 (94.4%)
# Gaps:           1/503 ( 0.2%)
# Score: 2427
#
#
#=======================================

CBP                1 LGYCCGRKYEFSPQTLCCYGKQLCTIPRDAAYYSYQNRYHFCEKCFTEIQ     50
                     ||||||||.|||||||||||||||||||||.|||||||||||||||.|||
p300               1 LGYCCGRKLEFSPQTLCCYGKQLCTIPRDATYYSYQNRYHFCEKCFNEIQ     50

CBP               51 GENVTLGDDPSQPQTTISKDQFEKKKNDTLDPEPFVDCKECGRKMHQICV    100
                     ||:|:||||||||||||:|:||.|:||||||||.||:|.|||||||||||
p300              51 GESVSLGDDPSQPQTTINKEQFSKRKNDTLDPELFVECTECGRKMHQICV    100

CBP              101 LHYDIIWPSGFVCDNCLKKTGRPRKENKFSAKRLQTTRLGNHLEDRVNKF    150
                     ||::||||:|||||.||||:.|.|||||||||||.:||||..||:|||.|
p300             101 LHHEIIWPAGFVCDGCLKKSARTRKENKFSAKRLPSTRLGTFLENRVNDF    150

CBP              151 LRRQNHPEAGEVFVRVVASSDKTVEVKPGMKSRFVDSGEMSESFPYRTKA    200
                     ||||||||:|||.||||.:||||||||||||:||||||||:|||||||||
p300             151 LRRQNHPESGEVTVRVVHASDKTVEVKPGMKARFVDSGEMAESFPYRTKA    200

CBP              201 LFAFEEIDGVDVCFFGMHVQEYGSDCPPPNTRRVYISYLDSIHFFRPRCL    250
                     |||||||||||:||||||||||||||||||.||||||||||:|||||:||
p300             201 LFAFEEIDGVDLCFFGMHVQEYGSDCPPPNQRRVYISYLDSVHFFRPKCL    250
```

```
CBP                251 RTAVYHEILIGYLEYVKKLGYVTGHIWACPPSEGDDYIFHCHPPDQKIPK    300
                       ||||||||||||||||||||||.|||||||||||||||||||||||||||
p300               251 RTAVYHEILIGYLEYVKKLGYTTGHIWACPPSEGDDYIFHCHPPDQKIPK    300

CBP                301 PKRLQEWYKKMLDKAFAERIIHDYKDIFKQATEDRLTSAKELPYFEGDFW    350
                       |||||||||||||||.:|||:|||||||||||||||||||||||||||||
p300               301 PKRLQEWYKKMLDKAVSERIVHDYKDIFKQATEDRLTSAKELPYFEGDFW    350

CBP                351 PNVLEESIKELEQEEEERKKEESTAASETTEGSQGDSKNAKKKNNKKTNK    400
                       |||||||||||||||||||:||:|: :|:|:.::||||||||||||||:|
p300               351 PNVLEESIKELEQEEEERKREENTS-NESTDVTKGDSKNAKKKNNKKTSK    399

CBP                401 NKSSISRANKKKPSMPNVSNDLSQKLYATMEKHKEVFFVIHLHAGPVINT    450
                       ||||:||.|||||.||||||||||||||||||||||||||.|.|||..|:
p300               400 NKSSLSRGNKKKPGMPNVSNDLSQKLYATMEKHKEVFFVIRLIAGPAANS    449

CBP                451 LPPIVDPDPLLSCDLMDGRDAFLTLARDKHWEFSSLRRSKWSTLCMLVEL    500
                       ||||||||||:.||||||||||||||||||.|||||||::|||:||||||
p300               450 LPPIVDPDPLIPCDLMDGRDAFLTLARDKHLEFSSLRRAQWSTMCMLVEL    499

CBP                501 HTQ    503
                       |||
p300               500 HTQ    502


#---------------------------------------
#---------------------------------------
```

### CBP>p300 (CBP:FUS Interaction Domain (CFID) | CBP Residues 1892-2441)

```
########################################
# Program: matcher
# Rundate: Tue  2 Apr 2024 13:48:43
# Commandline: matcher
#    -auto
#    -stdout
#    -asequence emboss_matcher-I20240402-134828-0445-34170096-p1m.asequence
#    -bsequence emboss_matcher-I20240402-134828-0445-34170096-p1m.bsequence
#    -datafile EBLOSUM62
#    -gapopen 14
#    -gapextend 4
#    -alternatives 1
#    -aformat3 pair
#    -sprotein1
#    -sprotein2
# Align_format: pair
# Report_file: stdout
########################################

#=======================================
#
# Aligned_sequences: 2
# 1: CBP_CFID
# 2: p300_CFID
# Matrix: EBLOSUM62
# Gap_penalty: 14
# Extend_penalty: 4
#
# Length: 553
# Identity:     286/553 (51.7%)
# Similarity:   344/553 (62.2%)
# Gaps:          65/553 (11.8%)
# Score: 988
#
#
#=======================================

CBP_CFID           1 TPT-QQPSTPQTPQPPAQPQPSPVSMSPAGFPSVARTQPPTTVSTGKPTS     49
                     ||| |||:|||||||.:||||:|.:..|   |.:.|||....||.||...
p300_CFID          1 TPTGQQPTTPQTPQPTSQPQPTPPNSMP---PYLPRTQAAGPVSQGKAAG     47

CBP_CFID          50 QV--PAPPPPAQPP-----PAAVEAARQIEREAQQQQHLYRVNINNSMPP     92
                     ||  |.||..||||     |||||.|.||:|.|:.|:.:..|.|...  |
p300_CFID         48 QVTPPTPPQTAQPPLPGPPPAAVEMAMQIQRAAETQRQMAHVQIFQR--P     95

CBP_CFID          93 GRTGMGTPGSQMAPVSLNVPRPNQ-VSGPVMPSMPPGQWQQAP------L    135
                     .:..| .|.:.|||:.:|.|...: .||.:.|.|.|...||.|      |
p300_CFID         96 IQHQM-PPMTPMAPMGMNPPPMTRGPSGHLEPGMGPTGMQQQPPWSQGGL    144

CBP_CFID         136 PQ-QQPMPGLPRPVISMQAQ-------AAVAGPRMPSVQP--PRSISPSA    175
                     || ||...|:|||.:...||       |...|.....:.|  |.::|..|
p300_CFID        145 PQPQQLQSGMPRPAMMSVAQHGQPLNMAPQPGLGQVGISPLKPGTVSQQA    194

CBP_CFID         176 LQDLLRTLKSPSSPQQQQQVLNILKSNPQLMAAFIKQRTAKYVANQPGMQ    225
                     ||:|||||:|||||.||||||:||.:||||:|||||||.|||..:.|...
p300_CFID        195 LQNLLRTLRSPSSPLQQQQVLSILHANPQLLAAFIKQRAAKYANSNPQPI    244
```

```
CBP_CFID         226 P-QPGL-QSQPGMQP-----QPGMHQQPSLQNLNAMQAGVPRPGVPPQQQ    268
                     | |||: |.|||:||     |.|:|..|::||:|.|||||.|.|:| |||
p300_CFID        245 PGQPGMPQGQPGLQPPTMPGQQGVHSNPAMQNMNPMQAGVQRAGLP-QQQ    293

CBP_CFID         269 AMGGLNPQGQALNIMNPGHNPNMASMNPQYREMLRRQLLQQQQQQQQQQQ    318
                     ....|.|....::......|.|..:|..|:|::||||          |..
p300_CFID        294 PQQQLQPPMGGMSPQAQQMNMNHNTMPSQFRDILRRQ----------QMM    333

CBP_CFID         319 QQQQQQQGSAGMAGGMAGHGQFQQPQGPGGYPPAMQQQQRMQQHLPLQGS    368
                     ||||||....|:..|||.|.|||||||.| |||  |||||||.|:  |..
p300_CFID        334 QQQQQQGAGPGIGPGMANHNQFQQPQGVG-YPP--QQQQRMQHHM--QQM    378

CBP_CFID         369 SMGQMAAQMGQLGQMGQPGLGADSTPNIQQALQQRILQQQQMKQQIGSPG    418
                     ..|.|    ||:||:.| .|||::..::| |.|||:||||     :|||.
p300_CFID        379 QQGNM----GQIGQLPQ-ALGAEAGASLQ-AYQQRLLQQQ-----MGSPV    417

CBP_CFID         419 QPNPMSPQQHMLSGQPQASHLPGQQIATSLSNQVRSPAPVQSPRPQSQPP    468
                     ||||||||||||..|.|:.||.||||..|||||||||.||.|||||||||
p300_CFID        418 QPNPMSPQQHMLPNQAQSPHLQGQQIPNSLSNQVRSPQPVPSPRPQSQPP    467

CBP_CFID         469 HSSPSPRIQPQPSPHHVSPQTGSPHPGLAVTMASSIDQGHLGNPEQSAML    518
                     |||||||:|||||||||||||.||||||....|:.::|||..:|:|::||
p300_CFID        468 HSSPSPRMQPQPSPHHVSPQTSSPHPGLVAAQANPMEQGHFASPDQNSML    517

CBP_CFID         519 PQL    521
                     .||
p300_CFID        518 SQL    520


#---------------------------------------
#---------------------------------------
```

### CBP>p300 (IDR6 | CBP Residues 1856-2090)

```
########################################
# Program: matcher
# Rundate: Tue  2 Apr 2024 17:59:58
# Commandline: matcher
#    -auto
#    -stdout
#    -asequence emboss_matcher-I20240402-175954-0242-92798871-p1m.asequence
#    -bsequence emboss_matcher-I20240402-175954-0242-92798871-p1m.bsequence
#    -datafile EBLOSUM62
#    -gapopen 14
#    -gapextend 4
#    -alternatives 1
#    -aformat3 pair
#    -sprotein1
#    -sprotein2
# Align_format: pair
# Report_file: stdout
########################################

#=======================================
#
# Aligned_sequences: 2
# 1: EMBOSS_001
# 2: EMBOSS_001
# Matrix: EBLOSUM62
# Gap_penalty: 14
# Extend_penalty: 4
#
# Length: 262
# Identity:     126/262 (48.1%)
# Similarity:   152/262 (58.0%)
# Gaps:          33/262 (12.6%)
# Score: 362
#
#
#=======================================

EMBOSS_001         1 QHRLQQAQLMRRRMATMNTRNV--PQQSLPSPTSAPPGTPT-QQPSTPQT     47
                     ||||||||::|||||:|....|  .||.|||||.|.|.||| |||:||||
EMBOSS_001         1 QHRLQQAQMLRRRMASMQRTGVVGQQQGLPSPTPATPTTPTGQQPTTPQT     50

EMBOSS_001        48 PQPPAQPQPSPVSMSPAGFPSVARTQPPTTVSTGKPTSQV--PAPPPPAQ     95
                     |||.:||||:|.:..|   |.:.|||....||.||...||  |.||..||
EMBOSS_001        51 PQPTSQPQPTPPNSMP---PYLPRTQAAGPVSQGKAAGQVTPPTPPQTAQ     97

EMBOSS_001        96 PP-----PAAVEAARQIEREAQQQHLYRVNINNSMPPGRTGMGTPGSQM    140
                     ||     |||||.|.||:|.|:.|:.:..|.|  ...|.:..| .|.:.|
EMBOSS_001        98 PPLPGPPPAAVEMAMQIQRAAETQRQMAHVQI--FQRPIQHQM-PPMTPM    144

EMBOSS_001       141 APVSLNVPRPNQ-VSGPVMPSMPPGQWQQAP------LPQ-QQPMPGLPR    182
                     ||:.:|.|...: .||.:.|.|.|...||.|      ||| ||...|:||
EMBOSS_001       145 APMGMNPPPMTRGPSGHLEPGMGPTGMQQQPPWSQGGLPQPQQLQSGMPR    194

EMBOSS_001       183 PVISMQAQ-------AAVAGPRMPSVQP--PRSISPSALQDLLRTLKSPS    223
                     |.:...||       |...|.....:.|  |.::|..|||:|||||:|||
EMBOSS_001       195 PAMMSVAQHGQPLNMAPQPGLGQVGISPLKPGTVSQQALQNLLRTLRSPS    244
```

```
EMBOSS_001       224 SPQQQQVLNIL    235
                     ||.||||||:||
EMBOSS_001       245 SPLQQQVLSIL    256


#---------------------------------------
#---------------------------------------
```

### CBP>p300 (IDR7 | CBP Residues 2109-2434)

```
########################################
# Program: matcher
# Rundate: Tue  2 Apr 2024 18:03:45
# Commandline: matcher
#    -auto
#    -stdout
#    -asequence emboss_matcher-I20240402-180332-0958-38107740-p1m.asequence
#    -bsequence emboss_matcher-I20240402-180332-0958-38107740-p1m.bsequence
#    -datafile EBLOSUM62
#    -gapopen 14
#    -gapextend 4
#    -alternatives 1
#    -aformat3 pair
#    -sprotein1
#    -sprotein2
# Align_format: pair
# Report_file: stdout
########################################

#=======================================
#
# Aligned_sequences: 2
# 1: EMBOSS_001
# 2: EMBOSS_001
# Matrix: EBLOSUM62
# Gap_penalty: 14
# Extend_penalty: 4
#
# Length: 305
# Identity:     171/305 (56.1%)
# Similarity:   202/305 (66.2%)
# Gaps:          29/305 ( 9.5%)
# Score: 683
#
#
#=======================================

EMBOSS_001         1 VANQPGM-QPQPGLQSQPGMQPQPGMHQQPSLQNLNAMQAGVPRPGVPPQ     49
                     :..|||| |.|||||. |.|..|.|:|..|::||:|.|||||.|.|:| |
EMBOSS_001         8 IPGQPGMPQGQPGLQP-PTMPGQQGVHSNPAMQNMNPMQAGVQRAGLP-Q     55

EMBOSS_001        50 QQAMGGLNPQGQALNIMNPGHNPNMASMNPQYREMLRRQLLQQQQQQQQQ     99
                     ||....|.|....:.......|.|..:|..|:|::||||          |
EMBOSS_001        56 QQPQQQLQPPMGGMSPQAQQMNMNHNTMPSQFRDILRRQ----------Q     95

EMBOSS_001       100 QQQQQQQQQGSAGMAGGMAGHGQFQQPQGPGGYPPAMQQQQRMQQHLPLQ    149
                     ..||||||....|:..|||.|.|||||||.| |||  |||||||.|:
EMBOSS_001        96 MMQQQQQQGAGPGIGPGMANHNQFQQPQGVG-YPP--QQQQRMQHHM---    139

EMBOSS_001       150 GSSMGQMAAQMGQLGQMGQPGLGADSTPNIQQALQQRILQQQQMKQQIGS    199
                      ..|.|  ..|||:||:.| .|||::..::| |.|||:||||     :||
EMBOSS_001       140 -QQMQQ--GNMGQIGQLPQ-ALGAEAGASLQ-AYQQRLLQQQ-----MGS    179

EMBOSS_001       200 PGQPNPMSPQQHMLSGQPQASHLPGQQIATSLSNQVRSPAPVQSPRPQSQ    249
                     |.||||||||||||..|.|:.||.||||..|||||||||.||.|||||||
EMBOSS_001       180 PVQPNPMSPQQHMLPNQAQSPHLQGQQIPNSLSNQVRSPQPVPSPRPQSQ    229
```

```
EMBOSS_001       250 PPHSSPSPRIQPQPSPHHVSPQTGSPHPGLAVTMASSIDQGHLGNPEQSA    299
                     |||||||||:|||||||||||||.||||||....|:.::|||..:|:|::
EMBOSS_001       230 PPHSSPSPRMQPQPSPHHVSPQTSSPHPGLVAAQANPMEQGHFASPDQNS    279

EMBOSS_001       300 MLPQL    304
                     ||.||
EMBOSS_001       280 MLSQL    284


#---------------------------------------
#---------------------------------------
```
